# Loss of excitatory inputs and decreased tonic and evoked activity of locus coeruleus neurons in aged P301S mice

**DOI:** 10.1101/2025.01.17.633373

**Authors:** Anthony M. Downs, Gracianne Kmiec, Christina M. Catavero, Zoé A. McElligott

## Abstract

Tau pathology in the locus coeruleus (LC) is associated with several neurodegenerative conditions including Alzheimer’s disease and frontotemporal dementia. Phosphorylated tau accumulates in the LC and results in inflammation, synaptic loss, and eventually cell death as the disease progresses. Loss of LC neurons and noradrenergic innervation is thought to contribute to the symptoms of cognitive decline later in disease. While loss and degeneration of LC neurons has been well studied, less is known about changes in LC physiology at advanced stages of tau pathology that precedes neurodegeneration. In this study, we investigated the *ex vivo* electrophysiological properties of LC neurons in male and female mice from the P301S mouse model of tauopathy at 9 months of age, a time-point when significant tau accumulation, cell death, and cognitive impairments are observed. We found a reduction in excitatory inputs and changes in excitatory post-synaptic current kinetics in male and female P301S. There was also a decrease in spontaneous discharge of LC neurons and an increase in AP threshold in P301S mice of both sexes. Finally, we observed a decrease in excitability and increase in rheobase current in P301S mice. Despite the decrease in LC activity in slice, we did not identify differences in total tissue norepinephrine (NE) or NE metabolites in prefrontal cortex or hippocampus. Together these findings demonstrate reductions in the activity and excitability of LC neurons at late stages of tau accumulation. However, compensatory mechanisms may maintain normal NE levels in LC projection regions *in* vivo.

## Introduction

The locus coeruleus (LC) is a dense collection of noradrenergic neurons located in the pons that innervates many cortical and subcortical structures throughout the brain (Robertson *et al*., 2013). Due to its broad innervation pattern, the LC is critical for a diverse array of functions including arousal, sleep-wake cycles, stress responses, learning and memory, and cognition (Mason, 1981; Harley, 1987; Kayama & Koyama, 2003; Aston-Jones & Cohen, 2005). The LC has been identified as a major site of phosphorylated tau accumulation during early Alzheimer’s Disease (AD) and results in progressive loss of LC neurons (Bondareff, Mountjoy & Roth, 1982; Zweig *et al*., 1988; Braak & Braak, 1991; Rüb *et al*., 2001; Andrés-Benito *et al*., 2017; Beardmore *et al*., 2024; Bueichekú *et al*., 2024). Loss of LC neurons and NE innervation is associated with cognitive decline in later stage AD, which suggests reduced LC-NE neurotransmission during AD progression (Jacobs *et al*., 2021).

While ample evidence links the LC to tau accumulation, there is relatively little known regarding the effects of tau pathology on LC physiology and connectivity itself. Generally speaking, genetic manipulation of tau has mixed effects on neurons. Tau knockout mice show reduced action potential firing in excitatory neurons in the somatosensory cortex but increased excitability of parvalbumin-positive inhibitory interneurons (Chang *et al*., 2021). Tau pathology is associated with cortical hypoactivity in a mouse model of combined β-amyloid (Aβ) and tau pathology (Busche *et al*., 2019). However, other studies have reported tau pathology-mediated hyperactivity in both mice and humans using PET imaging (Huijbers *et al*., 2019; Shimojo *et al*., 2020). A recent study, using the TgF344-AD transgenic rat model found hypoactivity of LC neurons at baseline relative to controls, but age-dependent changes in foot shock-evoked LC burst firing (Kelberman *et al*., 2023). Importantly, few of these studies have investigated sex differences in tau pathology induced changes in LC function. Our previous study investigating effects of alcohol exposure on the P301S genotype at 7 months of age identified effects of alcohol and sex on LC electrophysiological properties, but relatively few effects due to P301S genotype (Downs *et al*., 2023). Given the importance of the LC on a variety of cognitive and affective functions and its importance in the progression of AD and other tauopathies, further research is critically needed to understand LC dysfunction in aged mice, when tauopathy is expected to be most severe.

To investigate the effects of tau pathology on LC neuron physiology, we recorded LC neurons from the P301S (PS19) mouse model using *ex vivo* patch-clamp electrophysiology. P301S mice express the P301S variant of the human *MAPT* gene and show progressive synapse loss and tau accumulation with marked neuronal loss and cortical and hippocampal atrophy by 9 months of age (Yoshiyama *et al*., 2007). These mice also demonstrate behavioral phenotypes consistent with AD, including cognitive impairment and enhanced anxiety-like behaviors during aging, many of which are influenced by LC activity (Chalermpalanupap *et al*., 2018; Catavero *et al*., 2022). We found decreased excitatory neurotransmission to LC neurons in both male and female P301S mice at 9 months of age. We also found decreased spontaneous AP firing rate and decreased excitability in male and female P301S mice at 9 months of age. Despite the decrease in LC activity in slice, we did not identify differences in total tissue norepinephrine (NE) or NE metabolites in prefrontal cortex or hippocampus.

## Materials and Methods

### Animals

Adult male and female P301S (Jackson Labs #008160) mice on a C57BL/6J background and wild-type littermate controls were bred in-house. All mice were group housed and maintained on a standard 12/12-hr light cycle. Animals had *ad libitum* access to food and water. All animal procedures were performed in accordance with the regulations of the University of North Carolina at Chapel Hill’s institutional animal care and use committee. All mice used in this study were approximately 9-months old (39-42 weeks old) as there is evidence for significant tau accumulation and neuronal loss at this time point, but there is greater mortality at later time points (Yoshiyama *et al*., 2007).

### Immunohistochemistry

Brains were sliced coronally at 30 μm using a Leica CM3050S cryostat (Leica, Wetzlar, Germany) and stored in cryoprotectant at -20 °C. Slices were washed with 0.1 M phosphate buffer saline (PBS), pH 7.3, (2 x 10 minutes) and permeabilized in a solution of 0.5% bovine serum albumin (BSA) and 0.4% Triton-X100 in 0.1 M PBS for 10 minutes. Slices were then covered for 2 hours at room temperature with blocking buffer: 5.0% donkey serum in the 1% BSA and 0.4% Triton-X100 in PBS at pH 7.3. Slices were transferred to blocking buffer with primary antibody solution: rabbit anti-tyrosine hydroxylase at 1:1000 (Pel Freez, Rogers, United States) and mouse anti-human phospho-Tau (Ser 202, Thr 205) AT8 at 1:500 (Invitrogen, Waltham, United States) and incubated for 16 hours at 4°C with gentle shaking. After incubation, slices were washed with 0.1 M PBS (4 x 10 minutes) and a solution of 0.5% BSA and 0.4% Triton-X100 in 0.1 M PBS for 10 minutes. Slices were covered in blocking buffer for 1 hour and immediately transferred to blocking buffer with secondary antibodies: donkey anti-rabbit IgG AlexaFluor 555 (1:200; Invitrogen) and donkey anti-mouse IgG AlexaFluor 488 (1:200; Invitrogen) for 2 hours with gentle shaking while shielded from light. Subsequently, slices were rinsed with 0.1 M PBS (4 x 10 minutes). Slices were washed in 0.1 M PBS with DAPI at 1:1000 (Thermo Fisher Scientific, Waltham, United States) for 5 minutes. Slices were then mounted on charged slides and cover slipped with Vectashield Vibrance Antifade Mounting Medium (Vector Laboratories, Newark, California, United States).

### Image acquisition

Slices were imaged using a BZ-X800 Keyence (Keyence Corporation, Osaka, Japan) with a 20x objective. Imaging parameters and exposure settings were consistent among all images.

### Brain Slice preparation

Mice were deeply anesthetized with isoflurane, decapitated, and brains were removed and placed into ice-cold sucrose aCSF [in mM: 194 sucrose, 20 NaCl, 4.4 KCl, 2 CaCl_2_, 1.2 NaH_2_PO_4_. 10 glucose, 26 NaHCO_3_] oxygenated with 95% O_2_ 5% CO_2_ for slicing. Brains were sliced coronally at 200-300 μm using a Leica VT1000 vibratome (Germany). Brain slices were then incubated in oxygenated NMDG aCSF recovery solution [in mM: 92 NMDG, 2.5 KCl, 0.5 CaCl_2_, 10 MgSO_4_, 1.25 NaH_2_PO_4_, 25 glucose, 30 NaHCO_3_, 20 HEPES, 2 thiourea, 5 Na-ascorbate, and 3 Na-pyruvate] held at 32 °C for 12 minutes. Slices were then transferred to a holding container with HEPES aCSF [in mM: 92 NaCl, 2.5 KCl, 1.25 NaH_2_PO_4_, 30 NaHCO_3_, 20 HEPES, 2 thiourea, 5 Na-ascorbate, and 3 Na-pyruvate, 2 CaCl_2_, and 2 MgSO_4_] at 32 °C for at least 45 minutes. Slices were then transferred to a recording chamber and perfused with oxygenated aCSF [in mM: 124 NaCl, 4.4 KCl, 2 CaCl_2_, 1.2 MgSO_4_. 1 NaH_2_PO_4_, 10 glucose, 26 NaHCO_3_] at 30 °C at a constant rate of 2 mL/min.

### Whole-cell recordings

Recording electrodes (2-4 MΩ) were pulled on a P-97 Micropipette Puller (Sutter Instruments). All recordings were conducted with potassium-gluconate intercellular recording solution [in mM: 135 gluconic acid potassium, 5 NaCl, 2 MgCl_2_, 10 HEPES, 0.6 EGTA, 4 Na_2_ATP, 0.4 Na_2_GTP). All signals were acquired using an Axon Multiclamp 700B (Molecular Devices Sunnyvale, CA). Input resistance, holding current, and access resistance were continuously monitored throughout the experiment. Cells in which access resistance changed greater than 20% were excluded from analysis. 1-3 cells were recorded from each animal for each set of experiments to avoid oversampling from 1 animal.

LC neurons were identified by their anatomical location in the slice, morphology, absence of I_h_ currents, presence of spontaneous action potentials, and linear response to hyperpolarizing currents (Williams *et al*., 1984; Paladini, Beckstead & Weinshenker, 2007). Spontaneous glutamate-mediated excitatory postsynaptic currents (sEPSCs) were acquired in voltage clamp at a -80 mV holding potential. Spontaneous action potentials were acquired in current-clamp mode with no current injected (I=0). Excitability studies were performed in current clamp mode with current injected to hold cells to a common membrane potential of -75 mV to account for inter-cell variability. Rheobase currents were determined with a series of current ramps (0-120 pA at 120 pA/s and 100-220 pA at 120 pA/s). Action potential firing curves were then assessed with step-wise current injection ranging from -100 to +400 pA at 20 pA intervals, and the number of action potentials at each current step were counted. The same cells were used for both current clamp and voltage clamp experiments.

### Tissue Monoamines

Mice were euthanized by isoflurane inhalation and brains were rapidly dissected, sliced, and frozen on dry ice. Tissue punches containing the PFC and Hippocampus were collected and stored at -80 °C. For high performance liquid chromatography (HPLC) analysis of monoamines, tissue was homogenized in 100 mM perchloric acid by probe sonication at 4 °C and centrifuged at 10,000 g for 10 min. The supernatant was then filtered through a 0.45 µm PVDF membrane before HPLC. The cell pellet was retained, and the protein concentration was determined using a BCA assay (ThermoFisher, Waltham, MA, USA). Monoamines were then measured using HPLC with electrochemical detection. The HPLC system included an ESAS MD-150 x 3.2 mm column, an ESA 5020 guard cell, and an ESA 5600A Coularray detector with an ESA 6210 detector cell (ESA, Bradford, MA, USA). The guard cell potential was 475 mV, and the analytical cell potentials were 175, 100, 350, and 450 mV. A flow rate of 0.4 mL/min was used and the mobile phase contained [in mM], [1.7] 1-octanesulfonic acid sodium, [75] NaH_2_PO_4_, 0.25% triethylamine, and 8% acetonitrile at pH 2.9. Both retention time and electrochemical profile were used to identify monoamines, and their properties were compared to known standards.

### Data analysis

All electrophysiological data were analyzed using pClamp 10.6 software (Molecular Devices) and Easy Electrophysiology. All data were analyzed using a two way or three-way regular or repeated-measures ANOVA depending on the number of variables assessed in each experiment. The Greenhouse-Geisser sphericity correction was used for all 2-way ANOVAS. Significant main or interaction effects were subsequently analyzed using Tukey’s multiple comparisons test to control for multiple comparisons. Due to known sex differences in the noradrenergic system, we a priori performed post-hoc analyses on significant main effects. All data are expressed as mean ± SEM. P values ≤ 0.05 were considered statistically significant. All statistical tests were performed using Graphpad Prism 10.2 (La Jolla, CA, USA).

## Results

### Tau accumulates in LC at 9-months-old in male and female P301S mice

Because the LC is an early site of phosphorylated tau accumulation in both humans and rodents (Bondareff, Mountjoy & Roth, 1982; Zweig *et al*., 1988; Braak & Braak, 1991; Andrés-Benito *et al*., 2017; Zhu *et al*., 2018; Beardmore *et al*., 2024; Bueichekú *et al*., 2024), we first assessed whether phosphorylated tau is present in LC neurons of P301S mice at 9 months of age. While it is known that different post translational modifications of tau can affect disease severity, we focused on the S202, T205 (AT8) phosphorylated tau. We did not observe any AT8 staining in either male or female WT animals (**Fig 1A & B**), with the exception of small amounts of background staining of what are presumably blood vessels. When we examined male and female P301S mice, we observed AT8 in the perikarya of both TH-positive and TH-negative neurons in the LC and peri-LC (**Fig 1C & D**). We did not observe AT8 in the surrounding Barrington’s nucleus or parabrachial nucleus. While we did not quantify AT8 expression in this experiment, we did not observe qualitative differences in the number of AT8+ cells between males and females.

**Fig 1.**
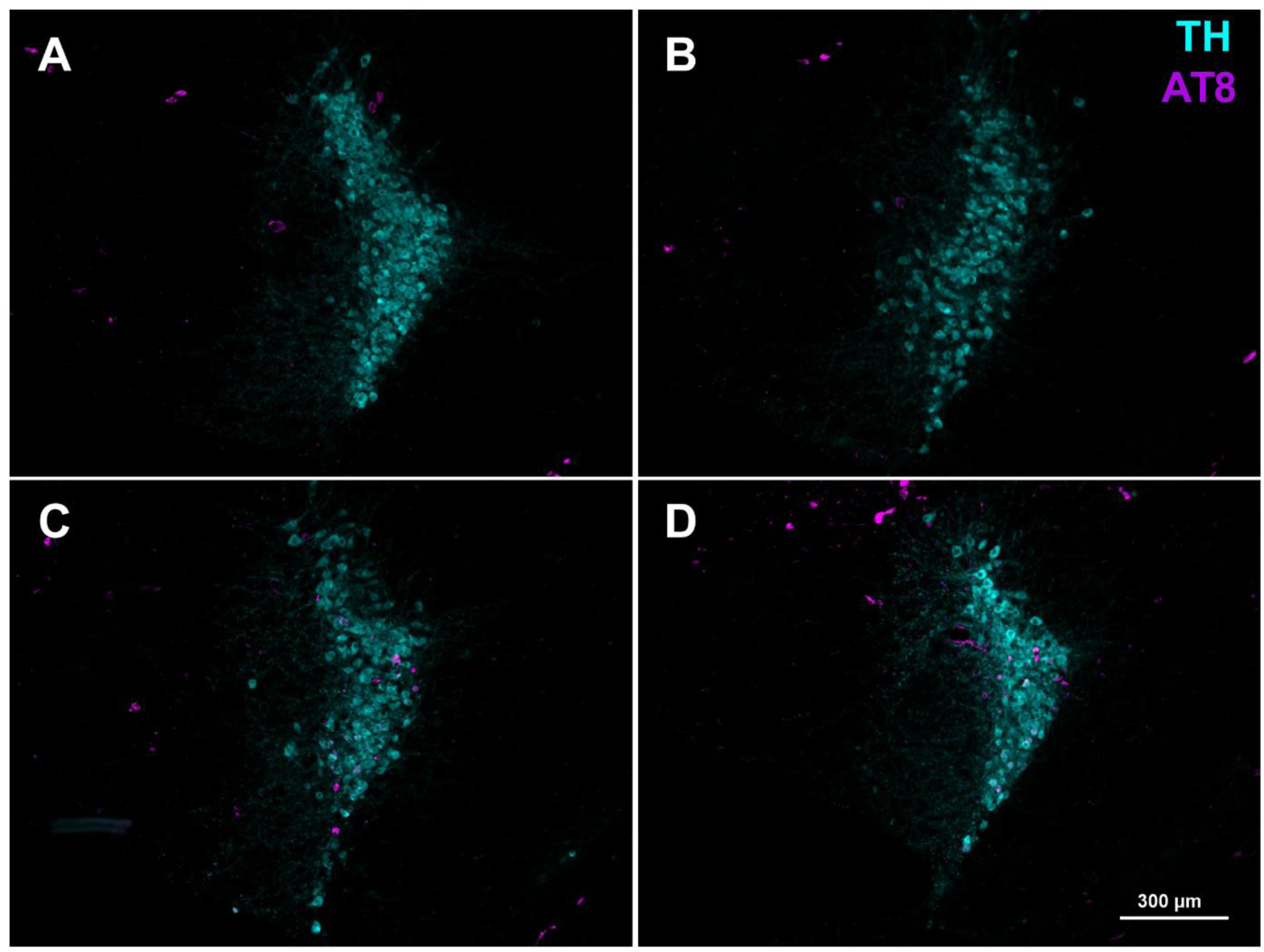
Phosphorylated tau (AT8) accumulates in the LC at 9 months of age. Representative images of WT Male (**A**), WT Female (**B**), P301S Male (**C**), P301S Female (**D**) LC neurons demonstrating that tau accumulates in both catecholaminergic and non-catecholaminergic neurons at 9 months of age. Tyrosine hydroxylase (TH) is shown in cyan and phosphorylated tau (AT8) is shown in magenta.

### Electrophysiological properties of LC neurons

We first assessed changes in resistance and capacitance in male and female WT and P301S mice in voltage clamp mode by applying steps from -70 mV to -80 mV. We did not observe any changes in whole cell resistance in P301S mice (main effect of genotype, F_(1,72)_= 0.179, p = 0.67) or between the sexes (main effect of sex, F_(1,72)_ = 3.231, p = 0.076) (**Table 1**). There were no significant changes in capacitance in P301S mice (main effect of genotype, F_(1,69)_ = 0.239, p = 0.626) or between the sexes (main effect of sex, F_(1,69)_ = 1.45, p = 0.232) (**Table 1**). There were no significant changes in access resistance by sex (main effect of sex, F_1,69_ = 0.71, p = 0.40) or genotype (main effect of genotype F_1,69_ = 0.32, p = 0.57) (**Table 1**).

**Table 1.**
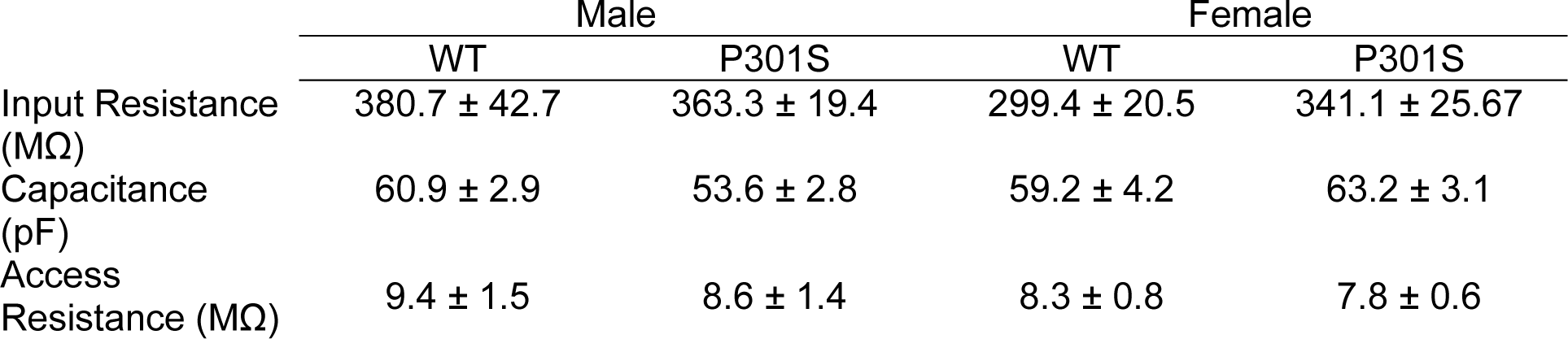
Membrane properties of locus coeruleus neurons. There were no significant differences by sex or P301S genotype. Data expressed as mean ± SEM.

### Spontaneous excitatory neurotransmission

We next assessed changes in glutamatergic transmission onto LC neurons by measuring spontaneous excitatory postsynaptic currents (sEPSCs) in the absence of tetrodotoxin in male and female P301S and WT littermates. We found a significant reduction in sEPSC frequency in P301S mice (main effect of genotype, F_(1,72)_ = 9.55, p = 0.003; Females: WT vs P301S, Tukey’s multiple comparisons test, p = 0.019) (**Fig 2A & B**). Male mice, regardless of genotype, also had a significantly lower sEPSC frequency in their LC neurons, as compared to female mice (main effect of sex, F_(1,72)_ = 7.00, p = 0.01). There were no significant effects on sEPSC amplitude due to P301S genotype or sex (**Fig 2C**). We found additional changes in the kinetics of sEPSCs. There was a significant reduction in decay time in P301S mice relative to WT (main effect of genotype, F_(1,72)_ = 5.77, p = 0.019) (**Fig 2D**). However, we did not observe any effects of sex on decay time (main effect of sex, F_(1,72)_ = 0.220, p = 0.640). Corresponding with the decrease in decay time, we also observed a significant decrease in AUC in P301S mice (main effect of genotype, F_(1,72)_ = 4.315, p = 0.041) (**Fig 2E**). Taken together these results demonstrate an overall reduction in excitatory neurotransmission in the LC of 9-month-old P301S animals.

**Figure 2.**
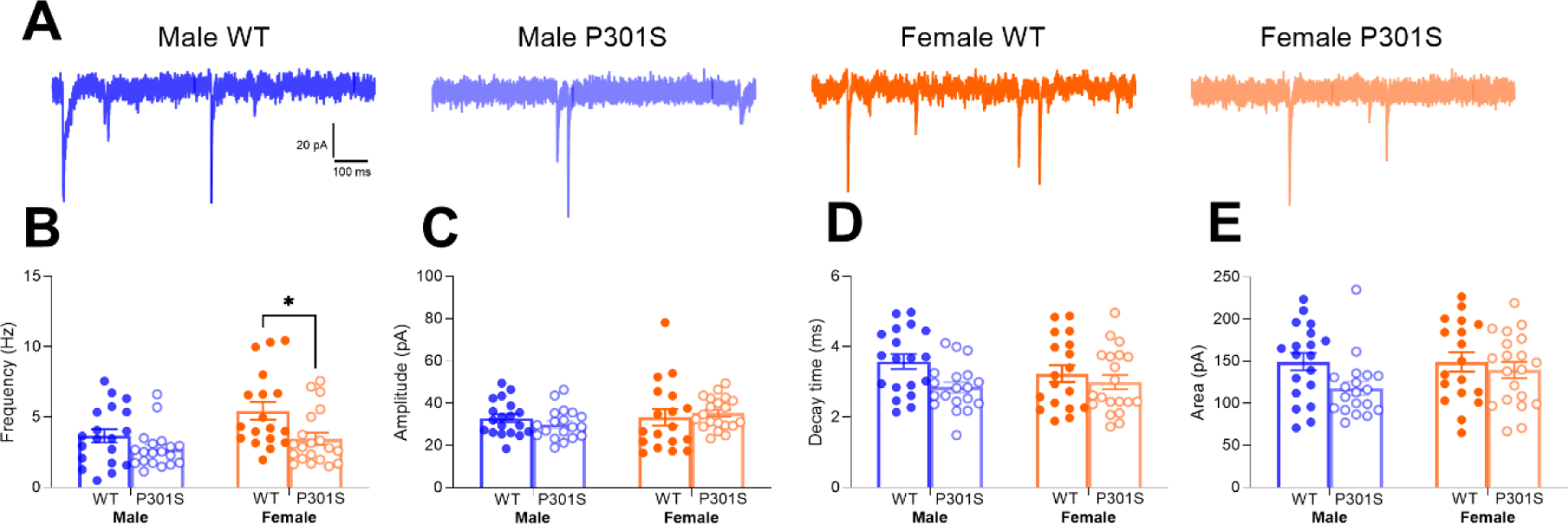
Spontaneous EPSCs in the LC are altered by tauopathy. (**A**) Representative traces from male and female P301S and WT LC neurons. **(B)** sEPSC frequency is reduced in male and female P301S mice **(C)** sEPSC amplitude is not altered by genotype or sex. **(D)** sEPSC decay time is reduced in male and female P301S mice. **(E)** sEPSC AUC is reduced in male and female P301S mice. Data expressed as mean ± S.E.M.

### Spontaneous action potentials

We next assessed spontaneous firing rate and action potential kinetics in WT and P301S mice. We found a significant reduction in AP firing frequency in P301S mice relative to WT (main effect of genotype, F_(1,71)_ = 13.24, p = 0.0005;, Tukey’s post hoc test, Female: P301S vs WT, p = 0.025) (**Fig 3A & B**). We did not observe any effects of sex on AP firing rate (main effect of sex, F_(1,71)_ = 0.011, p = 0.92). We did not observe any changes in AP half-width based on genotype, but there was a significant increase in AP half-width in female mice (main effect of sex, F_(1,71)_ = 4.85, p = 0.031) (**Fig 3C**). There were no significant changes in after-hyperpolarization potential (AHP) due to either genotype or sex (**Fig 3D**). However, there was a significant increase in AP threshold voltage in P301S mice (main effect of genotype, F_(1,71)_ = 6.16, p = 0.016) (**Fig 3E**).

**Figure 3.**
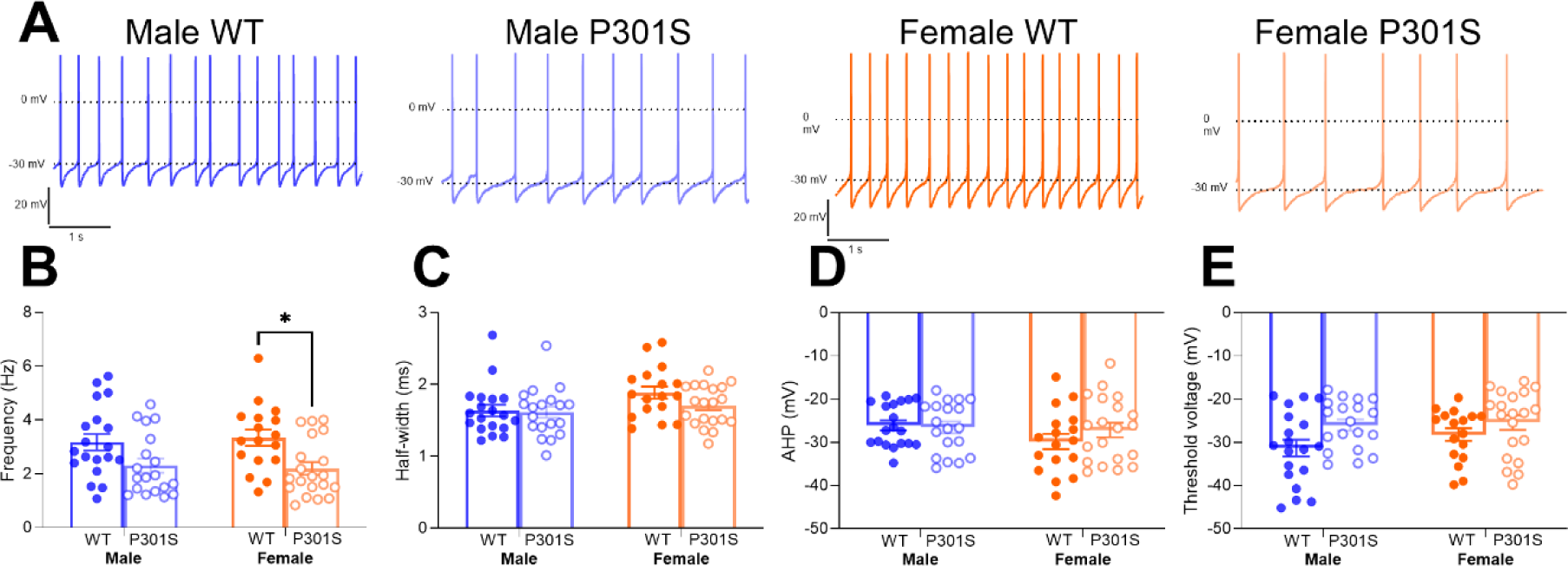
LC action potential frequency is reduced in P301S mice. **(A)** Representative traces form male and female P301S and WT LC neurons. **(B)** Spontaneous LC firing rate is reduced in male and female P301S mice. **(C)** Action-potential half-width is reduced in male mice but unaffected by genotype. **(D)** AHP is unaffected by sex or genotype. **(E)** AP threshold voltage is increased in male and female P301S mice. Data expressed as mean ± S.E.M.

### LC excitability

We next assessed the excitability of LC neurons. We first measured the rheobase current by first holding the cells at -75 mV to stop spontaneous firing and then applying ramps of increasing positive current injection. There was a significant increase in rheobase current in P301S mice relative to WT (main effect of genotype, F_(1,44)_ = 25.26, p < 0.0001; Males: WT vs P301S, Tukey’s post hoc test, p = 0.015; Females: WT vs P301S, Tukey’s post hoc test, p = 0.002) (**Fig 4B**). There was no effect of sex on rheobase current (main effect of sex, F_(1,44)_ = 0.719, p = 0.401). We next measured the number of evoked action potentials fired in response to positive current injection (0-400 pA, 20 pA steps). In male mice, increasing amounts of injected current increased the number action potentials fired per current step in both genotypes (main effect of current input, F_(1.444,33.21)_ = 112.8, p < 0.0001) (**Fig 4 A & C**). However, WT mice had a larger number of action potentials fired for each current step than P301S mice (current input x genotype interaction effect, F_(20,460)_ = 7.055, p < 0.0001; main effect of genotype, F_(1,23)_ = 11.89, p = 0.002). We observed similar result in female mice. Female WT mice had a larger number of action potentials fired for each current step than P301S females (current input x genotype interaction effect, F_(20,460)_ = 7.001, p < 0.0001; main effect of genotype, F_(1,23)_ = 8.487, p = 0.009) (**Fig 4 A & D**). We also compared the firing curves of each genotype type across sexes, but we did not observe any significant sex differences (three-way ANOVA, main effect of sex, F_(1,46)_ = 0.405, p = 0.528). Taken together, these results demonstrate a reduction in excitability of LC neurons in 9-month-old P301S mice.

**Figure 4.**
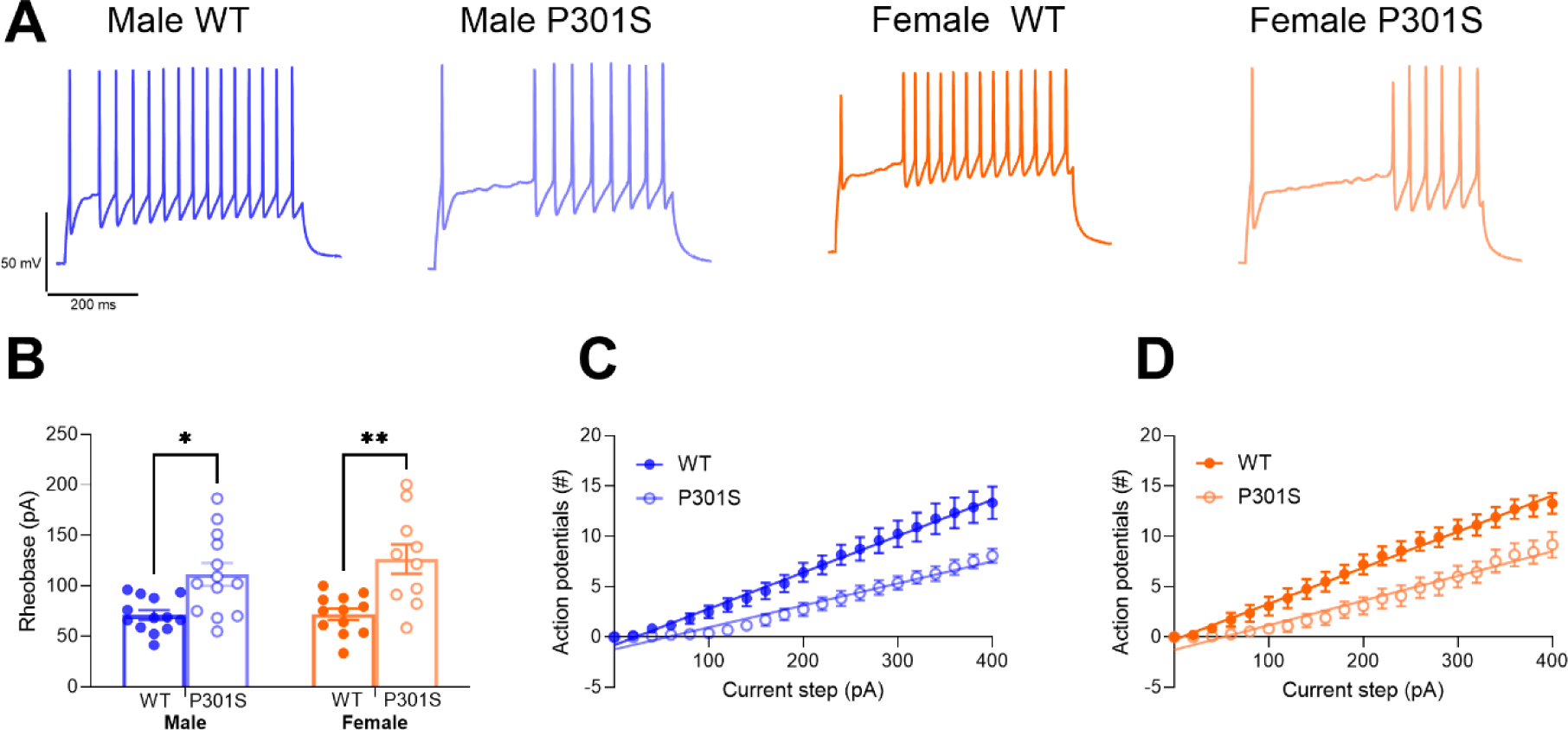
LC neurons are less excitability in P301S mice. **(A)** Representative traces of LC responses to 400 pA current injections. **(B)** Rheobase current is increased in male and female P301S mice. **(C)** LC neurons from male P301S mice are less excitable than LC neurons from WT male mice. **(D)** LC neurons from female P301S mice are less excitable than LC neurons from WT female mice. Data expressed as mean ± S.E.M.

### Tissue monoamine content in PFC and Hippocampus

We next investigated if there is a loss of NE and other catecholamine content in the PFC and hippocampus caused by either functional deficits in LC neurons or overt loss of LC innervation. We focused on the PFC and hippocampus because they receive NE innervation primarily from the LC, as opposed to other noradrenergic nuclei (Robertson *et al*., 2013). In the PFC, we found significantly more NE in females (main effect of sex, F_1,32_ = 10.41, p = 0.0029), but there were no differences based on genotype (main effect of genotype, F_1,32_ = 0.51, p = 0.48) (**Figure 5A**). We also found a trend towards more MHPG in females, but this was not significantly different (main effect of sex, F_1,27_ = 3.59, p = 0.069) (**Figure 5B**). There were no differences based on either genotype or sex in NE/MHPG ratio (**Figure 5C**). We also examined dopamine (DA) and metabolites in the PFC. Surprisingly, DA was differently altered based on genotype and sex. Male P301S mice had more DA than male WT mice, but DA was not altered based on genotype in female mice (genotype x sex interaction effect, F_1,32_ = 5.22, p = 0.29, Male WT vs Male P301S, Tukey’s multiple comparisons test, p = 0.015) (**Figure 5D**). We saw a similar sex and genotype interaction for the DA metabolite DOPAC (genotype x sex interaction effect, F_1,32_ = 6.44, p = 0.016, Male WT vs Male P301S, Tukey’s multiple comparisons test, p = 0.007) and a trend towards a similar interaction effect in HVA (genotype x sex interaction effect, F_1,32_ = 2.76, p = 0.11) (**Figure 5E & F**). The DA/DOPAC ratio (main effect of sex, F_1,32_ = 14.91, p = 0.0005) was increased in males but was unaltered by genotype (**Table 5G**). We also investigated the serotonin system in the PFC and found increased 5-HT (main effect of sex, F_1,32_ = 9.87, p = 0.004) and 5-HIAA (main effect of sex, F_1,27_ = 18.45, p = 0.0002) in females, but no differences based on genotype (**Figure 5H & I**).

**Figure 5.**
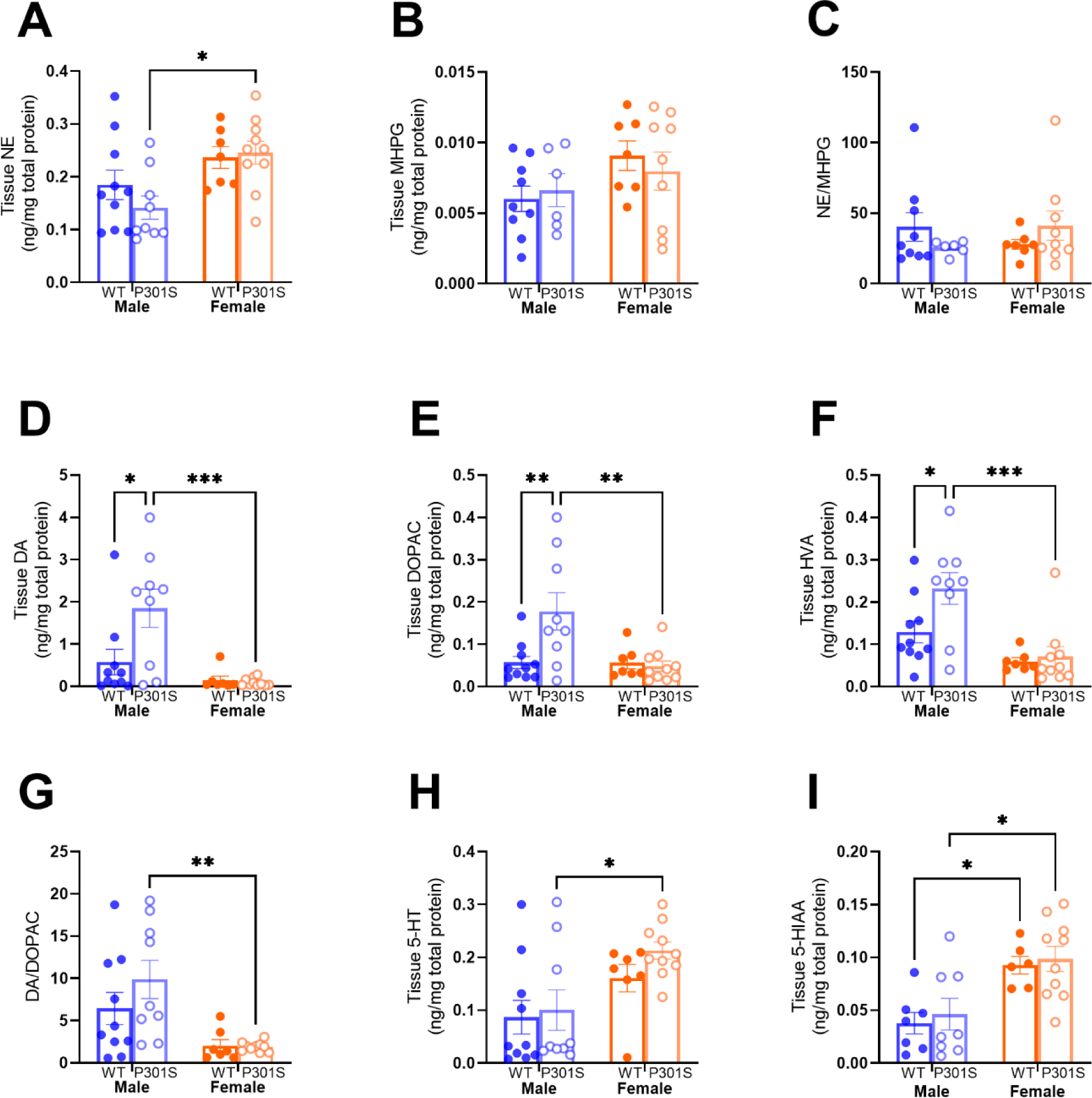
Tissue catecholamines in PFC. **(A)** NE content is reduced in Male P301S mice. **(B)** MHPG content is unchanged due to genotype or sex. **(C)** NE/MHPG ratio is unaltered. **(D)** DA content is increased in male P301S mice and reduced in female mice of both genotypes. **(E)** DOPAC content is increased in male P301S mice. **(F)** HVA content is increased in male P301S mice. **(G)** DA/DOAPC ratio is increased in male mice. **(H)**. 5-HT content is increased in female mice. **(I)** 5-HIAA content is increased in female mice.

In the hippocampus we did not find any significant differences in NE, MHPG, and NE/MHPG based on either genotype or sex (**Figure 6A-C**). We found a significant increase in DA in female mice (main effect of sex, F_1,32_ = 4.69, p = 0.038), but no effect based on genotype (main effect of genotype, F_1,32_ = 0.063, p = 0.43) (**Figure 6D**). We found a significant increase in DOPAC in female mice (main effect of sex, F_1,32_ = 25.19, p < 0.0001) and decreased DOPAC in female P301S mice versus female WT (main effect of genotype, F_1,32_ = 5.41, p = 0.027, Tukey’s multiple comparisons test, p = 0.032) (**Figure 6E**). We did not see any significant changes in HVA or DA/DOPAC ratio based on either genotype or sex (**Figure 6F & G**). We found a significant increase in both 5-HT (main effect of sex, F_1,32_ = 14.66, p = 0.0006) and 5-HIAA (main effect of sex, F_1,32_ = 25.84, p <0.0001) in female mice, but we did not observe changes based on genotype (**Figure 6H & I**).

**Figure 6.**
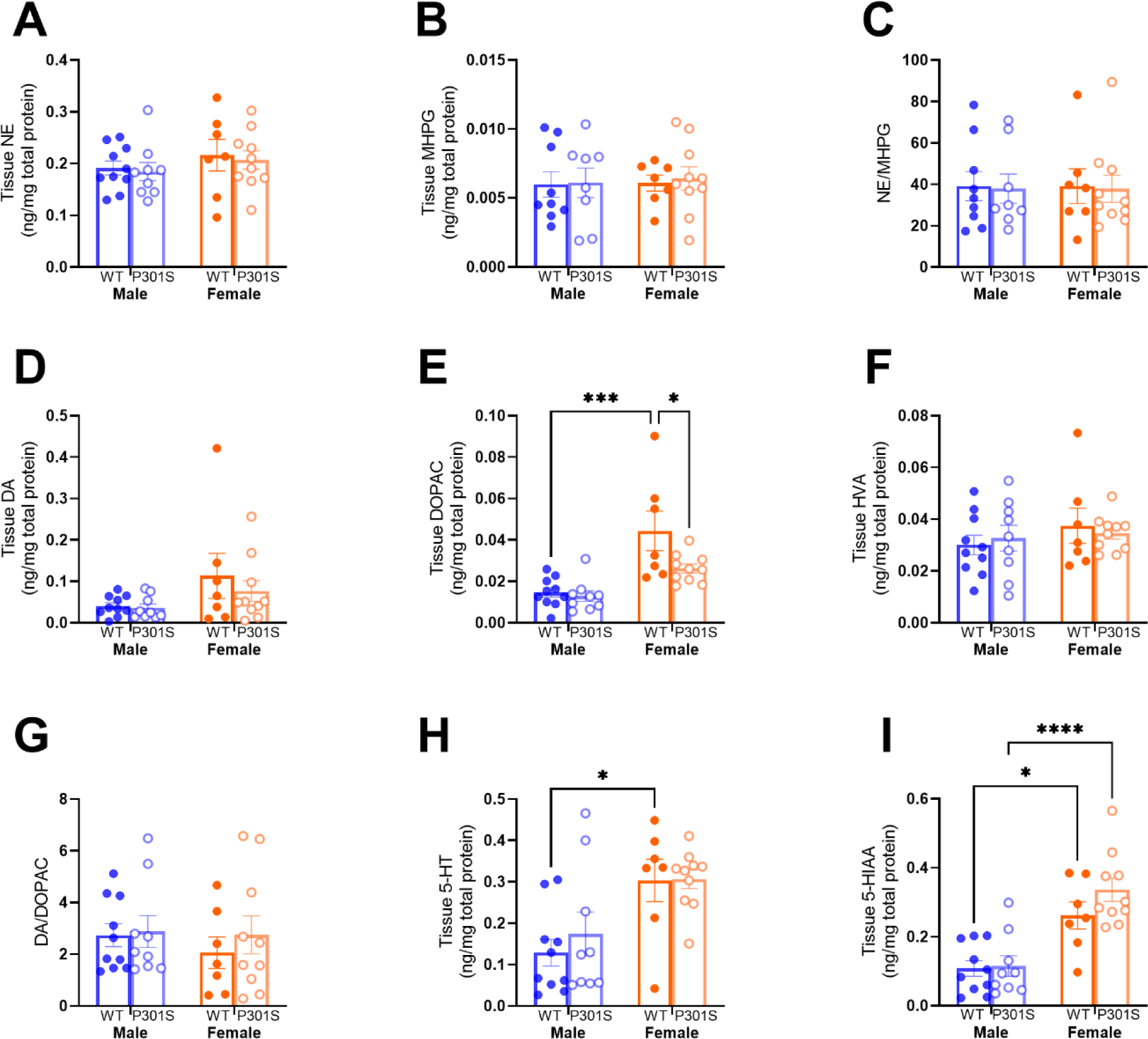
Tissue catecholamines in the hippocampus. **(A)** NE content, MHPG content **(B)**, and NE/MHPG ratio **(C)** were unaltered. (**D**) DA content is reduced in male mice. (**E**) DOPAC content is reduced in male mice and reduced in female P301S mice. (**F**) HVA content is unaltered. (**G**) DA/DOAPC ratio is unaltered. (**H**) 5-HT content is increased in female mice. (**I**) 5-HIAA content is increased in female mice.

## Discussion

In this study we examined the effects of tauopathy on LC physiology and plasticity in aged (9-month-old) male and female P301S mice and WT controls. At 9-months of age, we observed an increase in intercellular tau accumulation in noradrenergic LC neurons as well as accumulation in nearby non-noradrenergic cell populations. We found a significant reduction in glutamatergic inputs along with a decrease in charge transfer at AMPA receptors in male and female P301S mice. We also observed decreased tonic AP firing and an increase in threshold voltage of LC neurons in male and female P301S mice. Further, we observed decreased excitability following excitatory current injections in P301S mice of both sexes. However, many of these changes were more pronounced in female mice. Further, we did not find changes in NE or NE metabolites in the PFC or hippocampus based on genotype, but did observe changes in dopamine and serotonin.

At 9-months of age we observed a significant reduction in frequency and a reduction in both decay time and AUC of sEPSCs on LC neurons. This suggests both presynaptic changes in glutamatergic inputs and postsynaptic changes in AMPA subunit composition to reduce total charge transfer of excitatory inputs, as a reduction in frequency could be interpreted as both a loss of synapses or a reduction in presynaptic input. In our previous study examining LC physiology in 7-month-old P301S mice, we identified a decrease in sEPSC frequency and a decrease in sEPSC decay kinetics in male P301S mice, but we did not identify any changes in sEPSC properties in female P301S. This suggests that there may be sex differences in the time course of glutamatergic dysfunction in the LC in response to pathological tau accumulation (Downs *et al*., 2023). However, it is important to note that our previous study was conducted on singly housed animals on a reverse light-cycle, so the differences observed here may also reflect circadian activity and the environmental stressor of being singly housed. Exploring the relationship between sex, sleep/circadian period and LC function will be critical for future experiments. Indeed, the P301S mice show a dark-phase sleep disruption, and the onset of this disturbance is earlier in females. The same study also found that sleep disturbance promoted tau hyperphosphorylation in the LC (Martin *et al*., 2024). A number of previous studies have identified that pathological, phosphorylated tau disrupts post-synaptic density structure and impairs AMPA receptor insertion into the synapse (Gong & Lippa, 2010; Hoover *et al*., 2010; Jurado, 2017; Shrivastava *et al*., 2019). While studies using *ex vivo* electrophysiological recordings from tauopathy model mice are fairly limited, one recent study using the P301S mouse found a significant reduction in glutamatergic inputs to basal forebrain cholinergic neurons, and this effect was mediated by alterations in both AMPA and NMDA receptors (Zhong, Cao & Yan, 2022). One intriguing possibility is that, given that excess glutamatergic transmission is thought to be excitotoxic, the loss of glutamatergic inputs at both a presynaptic and postsynaptic level may be adaptive in the context of tauopathy and may protect remaining LC neurons from excitotoxicity (Smith, Gorospe & Kusiak, 2006; Vosler, Brennan & Chen, 2008; Dong, Wang & Qin, 2009; Supnet & Bezprozvanny, 2010). However, this likely fails at later stages of disease when overt LC neuron loss is observed (Chalermpalanupap *et al*., 2018). Future studies will need to assess the timeline of glutamatergic deficits in tauopathy and if reducing glutamatergic inputs is protective for LC neurons.

In the present study, we observed a decrease in the spontaneous firing rate of LC neurons in both male and female P301S mice at 9-months-old relative to WT controls. This contrasts with our previous study, where we did not find any change in the spontaneous firing rate of P301S LC neurons at 7-months-old, suggesting that LC dysfunction progresses with disease state (Downs *et al*., 2023). Notably, phosphorylated tau, particularly the AT-8 isoform is present in the LC of P301S mice in large quantities at 6-months of age, suggesting additional factors, such as synapse loss from excitatory inputs, may mediate the reduction in tonic firing rate (Kang *et al*., 2020). We also identified an increase in the threshold voltage of spontaneous APs in male and female P301S mice, which is consistent with the firing rate data. A recent *in vivo* electrophysiological study in the TgF344-AD rat model, which expresses the human disease-causing variants of amyloid precursor protein and presenilin-1, showed reduced tonic LC firing rates at both 6 months and 15 months of age (Kelberman *et al*., 2023). While AD-like pathology is driven by different transgenes in these rats, pre-tangle tau does accumulate in the LC of the TgF344-AD rat starting at 6 months (Rorabaugh *et al*., 2017). The fact that the appearance of pre-tangle tau coincides with the decrease in tonic LC firing rate in both TgF344-AD rats and our mice suggests that tau accumulation plays a role in the abnormal physiology of LC neurons, although additional insults, such as loss of excitatory inputs, likely contributes. Future studies to resolve how alterations in glutamatergic drive impact excitability will undoubtedly shed light on this physiology.

In addition to changes in firing rate, we also found LC neurons from P301S mice to be significantly less excitable in response to injection of positive current steps. Notably, in our previous study of 7-month-old P301S mice, we did not observe any significant changes in excitability, suggesting that LC function deteriorates with disease progression (Downs *et al*., 2023). As stated above, deeper analysis into how circadian period influences these phenomena will be undoubtedly informative. Current data on the effects of tau pathology on neuronal excitability are conflicting with some studies finding increased or decreased excitability in both hippocampal and cortical neuronal populations (Crimins, Rocher & Luebke, 2012; Busche *et al*., 2019; Huijbers *et al*., 2019; Shimojo *et al*., 2020; Brown *et al*., 2023). Studies using *MAPT* knockout mice have demonstrated contrasting effects in different neuronal populations, with *MAPT* knockout increasing excitability in inhibitory interneurons but having no effect in excitatory pyramidal cells (Chang *et al*., 2021). This suggests that the effects of tau on excitability are cell-type specific. The mechanisms leading to reduced excitability at 9 months of age are unclear. One previous study found that tau accumulation in the hippocampus reduced K_v_4.2 localization in dendrites (Hall *et al*., 2015). It’s possible that, in later stage tau pathology, tau may enhance K_v_4.2 accumulation in dendrites or alter other ion channels to reduce LC excitability. Alternatively, in addition to the decrease in excitatory inputs, tau may enhance inhibitory inputs to alter E/I balance and reduce firing rate and excitability. Our study did not block glutamatergic or GABAergic inputs when monitoring excitability so that we could examine the LC wholistically. However, future studies could mechanistically dissect if synaptic inputs influence the intrinsic excitability of LC neurons with and without tau pathology.

Our physiology data suggests that central NE-neurotransmission from the LC may be reduced in the later stages of tauopathy disorders. This contrasts with the early stages of AD and tauopathy disorders where NE neurotransmission is thought to be hyperactive and contribute to prodromal symptoms (Elrod *et al*., 1997; Ehrenberg *et al*., 2018; Weinshenker, 2018; Henjum *et al*., 2022). Ultimately, the loss of NE neurotransmission may worsen tauopathy progression, as previous studies have demonstrated exacerbated progression of neuropathology throughout the neuroaxis after ablation of LC neurons in P301S mice (Chalermpalanupap *et al*., 2018). The loss of coordinated NE neurotransmission may also contribute to the development of later stage symptoms of AD, such as apathy and neurocognitive impairments, which are associated with reduced LC activity (Matthews *et al*., 2002; Hezemans *et al*., 2022; Ye *et al*., 2022).

A loss of NE innervation and NE content in the hippocampus and forebrain areas has been identified in people and multiple mouse models of tauopathy and AD (Matthews *et al*., 2002; Francis *et al*., 2012; Rorabaugh *et al*., 2017; Chalermpalanupap *et al*., 2018). The functional changes in LC hypoactivity we observed suggest another mechanism by which forebrain NE content may be reduced. Surprisingly, we did not observe any changes in total tissue NE or the major NE metabolite MHPG in either PFC or hippocampus due to P301S genotype. This may reflect presynaptic adaptations that maintain tissue content of NE despite reduced activity of LC neurons *ex vivo*. Future studies could investigate release of NE in the PFC and hippocampus *in vivo* to determine if P301S mice have lower levels of NE release despite normal tissue content and explore if this release is dysregulated. Despite the lack of changes in NE, we did find altered DA and DA metabolite levels in the PFC, but not the hippocampus, in male P301S mice. Male P301S mice had increased DA and DA metabolites in this region, which may be of particular interest given the disruption of learning, cognitive flexibility, social behaviors that have been noted in various tauopathy models (Sydow *et al*., 2011; Chalermpalanupap *et al*., 2018; Samaey *et al*., 2019; Watt *et al*., 2020; Catavero *et al*., 2022). Future studies should investigate tau pathology in the VTA, a major source of PFC DA.

One limitation of our study is that we are investigating the entire, undifferentiated LC rather than investigating the LC in a module/efferent specific manner. Over the past decade, it has become clear that LC neurons send projections to various forebrain regions from topographically distinct regions of the LC along rostro/caudal and dorsal/ventral axes (Mason & Fibiger, 1979; Chandler, Gao & Waterhouse, 2014; Aston-Jones & Waterhouse, 2016). We did not investigate LC physiology in a topographically distinct way, which may obscure more aberrant physiology in more sensitive regions. For example, in early AD pre-tangle tau accumulation is worse in the middle third of the LC which provides dense innervation to the hippocampus and cortex (Waterhouse *et al*., 1983; Ehrenberg *et al*., 2017). Accordingly, LC physiology may have greater or different alterations in those regions relative to LC regions that are spared from tau accumulation. Future studies should investigate the topographical pattern of tau accumulation and emergent pathophysiology in the LC across tau pathology progression. This may provide key insights into mechanisms involved in the neuropsychiatric symptoms of AD and FTD across disease progression.

Taken together, our data demonstrate significant reductions in LC excitatory inputs, tonic firing rate, and excitability in both male in female P301S mice at 9 months of age. Given the broad innervation pattern and diverse roles of the LC in cognition, arousal, and stress responses, these physiological changes could have significant roles in the progression of tauopathy disorders. Future studies will be critical to understand the effects of tau pathology on LC physiology across the lifespan.

## Acknowledgements

This study was supported in part by the Emory HPLC Bioanalytical Core (EHBC), which is subsidized by the Emory University School of Medicine and is one of the Emory Integrated Core Facilities. Additional support was provided by the Georgia Clinical & Translational Science Alliance of the National Institutes of Health under Award Number UL1TR002378. The content is solely the responsibility of the authors and does not necessarily reflect the official views of the National Institutes of Health.

## Author Contributions

A.M.D designed and performed experiments, analyzed data, and wrote the manuscript. G.K designed and performed experiments, analyzed data, and wrote the manuscript. C.M.C designed and performed experiments and wrote the manuscript. Z.A.M. designed experiments, wrote the manuscript, and provided funding for the study. All authors approve the final manuscript and agree to be accountable for all aspects of the work.

## References

Andrés-Benito, P., Fernández-Dueñas, V., Carmona, M., Escobar, L.A., Torrejón-Escribano, B., Aso, E., Ciruela, F. & Ferrer, I. (2017) Locus coeruleus at asymptomatic early and middle Braak stages of neurofibrillary tangle pathology. Neuropathology and applied neurobiology, 43, 373–392.

Aston-Jones, G. & Cohen, J.D. (2005) An integrative theory of locus coeruleus-norepinephrine function: adaptive gain and optimal performance. Annual review of neuroscience, 28, 403–450.

Aston-Jones, G. & Waterhouse, B. (2016) Locus coeruleus: From global projection system to adaptive regulation of behavior. Brain research, 1645, 75–78.

Beardmore, R., Durkin, M., Zayee-Mellick, F., Lau, L.C., Nicoll, J.A.R., Holmes, C. & Boche, D. (2024) Changes in the locus coeruleus during the course of Alzheimer’s disease and their relationship to cortical pathology. Neuropathology and applied neurobiology, 50, e12965.

Bondareff, W., Mountjoy, C.Q. & Roth, M. (1982) Loss of neurons of origin of the adrenergic projection to cerebral cortex (nucleus locus ceruleus) in senile dementia. Neurology, 32, 164–168.

Braak, H. & Braak, E. (1991) Neuropathological stageing of Alzheimer-related changes. Acta neuropathologica, 82, 239–259.

Brown, J., Camporesi, E., Lantero-Rodriguez, J., Olsson, M., Wang, A., Medem, B., Zetterberg, H., Blennow, K., Karikari, T.K., Wall, M. & Hill, E. (2023) Tau in cerebrospinal fluid induces neuronal hyperexcitability and alters hippocampal theta oscillations. Acta Neuropathologica Communications, 11, 67.

Bueichekú, E., Diez, I., Kim, C.M., Becker, J.A., Koops, E.A., Kwong, K., Papp, K.V., Salat, D.H., Bennett, D.A., Rentz, D.M., Sperling, R.A., Johnson, K.A., Sepulcre, J. & Jacobs, H.I.L. (2024) Spatiotemporal patterns of locus coeruleus integrity predict cortical tau and cognition. Nat Aging, 4, 625–637.

Busche, M.A., Wegmann, S., Dujardin, S., Commins, C., Schiantarelli, J., Klickstein, N., Kamath, T.V., Carlson, G.A., Nelken, I. & Hyman, B.T. (2019) Tau impairs neural circuits, dominating amyloid-β effects, in Alzheimer models in vivo. Nature neuroscience, 22, 57–64.

Catavero, C.M., Marsh, A.E., Downs, A.M., Teklezghi, A.T., Cohen, T.J. & McElligott, Z.A. (2022) Effects of Long-Term Alcohol Consumption on Behavior in the P301S (Line PS19) Tauopathy Mouse Model. bioRxiv, 2022.2007.2012.499737.

Chalermpalanupap, T., Schroeder, J.P., Rorabaugh, J.M., Liles, L.C., Lah, J.J., Levey, A.I. & Weinshenker, D. (2018) Locus Coeruleus Ablation Exacerbates Cognitive Deficits, Neuropathology, and Lethality in P301S Tau Transgenic Mice. The Journal of neuroscience: the official journal of the Society for Neuroscience, 38, 74-92.

Chandler, D.J., Gao, W.-J. & Waterhouse, B.D. (2014) Heterogeneous organization of the locus coeruleus projections to prefrontal and motor cortices. Proceedings of the National Academy of Sciences, 111, 6816–6821.

Chang, C.W., Evans, M.D., Yu, X., Yu, G.Q. & Mucke, L. (2021) Tau reduction affects excitatory and inhibitory neurons differently, reduces excitation/inhibition ratios, and counteracts network hypersynchrony. Cell reports, 37, 109855.

Crimins, J.L., Rocher, A.B. & Luebke, J.I. (2012) Electrophysiological changes precede morphological changes to frontal cortical pyramidal neurons in the rTg4510 mouse model of progressive tauopathy. Acta neuropathologica, 124, 777–795.

Dong, X.X., Wang, Y. & Qin, Z.H. (2009) Molecular mechanisms of excitotoxicity and their relevance to pathogenesis of neurodegenerative diseases. Acta pharmacologica Sinica, 30, 379–387.

Downs, A.M., Catavero, C.M., Kasten, M.R. & McElligott, Z.A. (2023) Tauopathy and alcohol consumption interact to alter locus coeruleus excitatory transmission and excitability in male and female mice. *Alcohol (Fayetteville*, N.Y.), 107, 97–107.

Ehrenberg, A.J., Nguy, A.K., Theofilas, P., Dunlop, S., Suemoto, C.K., Di Lorenzo Alho, A.T., Leite, R.P., Diehl Rodriguez, R., Mejia, M.B., Rüb, U., Farfel, J.M., de Lucena Ferretti-Rebustini, R.E., Nascimento, C.F., Nitrini, R., Pasquallucci, C.A., Jacob-Filho, W., Miller, B., Seeley, W.W., Heinsen, H. & Grinberg, L.T. (2017) Quantifying the accretion of hyperphosphorylated tau in the locus coeruleus and dorsal raphe nucleus: the pathological building blocks of early Alzheimer’s disease. Neuropathology and applied neurobiology, 43, 393–408.

Ehrenberg, A.J., Suemoto, C.K., França Resende, E.P., Petersen, C., Leite, R.E.P., Rodriguez, R.D., Ferretti-Rebustini, R.E.L., You, M., Oh, J., Nitrini, R., Pasqualucci, C.A., Jacob-Filho, W., Kramer, J.H., Gatchel, J.R. & Grinberg, L.T. (2018) Neuropathologic Correlates of Psychiatric Symptoms in Alzheimer’s Disease. Journal of Alzheimer’s disease : JAD, 66, 115–126.

Elrod, R., Peskind, E.R., DiGiacomo, L., Brodkin, K.I., Veith, R.C. & Raskind, M.A. (1997) Effects of Alzheimer’s disease severity on cerebrospinal fluid norepinephrine concentration. The American journal of psychiatry, 154, 25–30.

Francis, B.M., Yang, J., Hajderi, E., Brown, M.E., Michalski, B., McLaurin, J., Fahnestock, M. & Mount, H.T. (2012) Reduced tissue levels of noradrenaline are associated with behavioral phenotypes of the TgCRND8 mouse model of Alzheimer’s disease. Neuropsychopharmacology : official publication of the American College of Neuropsychopharmacology, 37, 1934–1944.

Gong, Y. & Lippa, C.F. (2010) Review: disruption of the postsynaptic density in Alzheimer’s disease and other neurodegenerative dementias. American journal of Alzheimer’s disease and other dementias, 25, 547–555.

Hall, A.M., Throesch, B.T., Buckingham, S.C., Markwardt, S.J., Peng, Y., Wang, Q., Hoffman, D.A. & Roberson, E.D. (2015) Tau-dependent Kv4.2 depletion and dendritic hyperexcitability in a mouse model of Alzheimer’s disease. The Journal of neuroscience : the official journal of the Society for Neuroscience, 35, 6221–6230.

Harley, C.W. (1987) A role for norepinephrine in arousal, emotion and learning?: limbic modulation by norepinephrine and the Kety hypothesis. Progress in neuro-psychopharmacology & biological psychiatry, 11, 419–458.

Henjum, K., Watne, L.O., Godang, K., Halaas, N.B., Eldholm, R.S., Blennow, K., Zetterberg, H., Saltvedt, I., Bollerslev, J. & Knapskog, A.B. (2022) Cerebrospinal fluid catecholamines in Alzheimer’s disease patients with and without biological disease. Translational psychiatry, 12, 151.

Hezemans, F.H., Wolpe, N., O’Callaghan, C., Ye, R., Rua, C., Jones, P.S., Murley, A.G., Holland, N., Regenthal, R., Tsvetanov, K.A., Barker, R.A., Williams-Gray, C.H., Robbins, T.W., Passamonti, L. & Rowe, J.B. (2022) Noradrenergic deficits contribute to apathy in Parkinson’s disease through the precision of expected outcomes. PLoS computational biology, 18, e1010079.

Hoover, B.R., Reed, M.N., Su, J., Penrod, R.D., Kotilinek, L.A., Grant, M.K., Pitstick, R., Carlson, G.A., Lanier, L.M., Yuan, L.L., Ashe, K.H. & Liao, D. (2010) Tau mislocalization to dendritic spines mediates synaptic dysfunction independently of neurodegeneration. Neuron, 68, 1067–1081.

Huijbers, W., Schultz, A.P., Papp, K.V., LaPoint, M.R., Hanseeuw, B., Chhatwal, J.P., Hedden, T., Johnson, K.A. & Sperling, R.A. (2019) Tau Accumulation in Clinically Normal Older Adults Is Associated with Hippocampal Hyperactivity. The Journal of neuroscience : the official journal of the Society for Neuroscience, 39, 548–556.

Jacobs, H.I.L., Becker, J.A., Kwong, K., Engels-Domínguez, N., Prokopiou, P.C., Papp, K.V., Properzi, M., Hampton, O.L., d’Oleire Uquillas, F., Sanchez, J.S., Rentz, D.M., El Fakhri, G., Normandin, M.D., Price, J.C., Bennett, D.A., Sperling, R.A. & Johnson, K.A. (2021) In vivo and neuropathology data support locus coeruleus integrity as indicator of Alzheimer’s disease pathology and cognitive decline. Science translational medicine, 13, eabj2511.

Jurado, S. (2017) AMPA Receptor Trafficking in Natural and Pathological Aging. Frontiers in molecular neuroscience, 10, 446.

Kang, S.S., Liu, X., Ahn, E.H., Xiang, J., Manfredsson, F.P., Yang, X., Luo, H.R., Liles, L.C., Weinshenker, D. & Ye, K. (2020) Norepinephrine metabolite DOPEGAL activates AEP and pathological Tau aggregation in locus coeruleus. The Journal of clinical investigation, 130, 422–437.

Kayama, Y. & Koyama, Y. (2003) Control of sleep and wakefulness by brainstem monoaminergic and cholinergic neurons. Acta neurochirurgica. Supplement, 87, 3–6.

Kelberman, M.A., Rorabaugh, J.M., Anderson, C.R., Marriott, A., DePuy, S.D., Rasmussen, K., McCann, K.E., Weiss, J.M. & Weinshenker, D. (2023) Age-dependent dysregulation of locus coeruleus firing in a transgenic rat model of Alzheimer’s disease. Neurobiology of aging, 125, 98–108.

Luster, B.R., Cogan, E.S., Schmidt, K.T., Pati, D., Pina, M.M., Dange, K. & McElligott, Z.A. (2020) Inhibitory transmission in the bed nucleus of the stria terminalis in male and female mice following morphine withdrawal. Addiction biology, 25, e12748.

Martin, S.C., Joyce, K.K., Lord, J.S., Harper, K.M., Nikolova, V.D., Cohen, T.J., Moy, S.S. & Diering, G.H. (2024) Sleep Disruption Precedes Forebrain Synaptic Tau Burden and Contributes to Cognitive Decline in a Sex-Dependent Manner in the P301S Tau Transgenic Mouse Model. eNeuro, 11.

Mason, S.T. (1981) Noradrenaline in the brain: progress in theories of behavioural function. Prog Neurobiol, 16, 263–303.

Mason, S.T. & Fibiger, H.C. (1979) Regional topography within noradrenergic locus coeruleus as revealed by retrograde transport of horseradish peroxidase. The Journal of comparative neurology, 187, 703–724.

Matthews, K.L., Chen, C.P., Esiri, M.M., Keene, J., Minger, S.L. & Francis, P.T. (2002) Noradrenergic changes, aggressive behavior, and cognition in patients with dementia. Biological psychiatry, 51, 407–416.

Paladini, C.A., Beckstead, M.J. & Weinshenker, D. (2007) Electrophysiological properties of catecholaminergic neurons in the norepinephrine-deficient mouse. Neuroscience, 144, 1067–1074.

Robertson, S.D., Plummer, N.W., de Marchena, J. & Jensen, P. (2013) Developmental origins of central norepinephrine neuron diversity. Nature neuroscience, 16, 1016–1023.

Rorabaugh, J.M., Chalermpalanupap, T., Botz-Zapp, C.A., Fu, V.M., Lembeck, N.A., Cohen, R.M. & Weinshenker, D. (2017) Chemogenetic locus coeruleus activation restores reversal learning in a rat model of Alzheimer’s disease. Brain : a journal of neurology, 140, 3023–3038.

Rüb, U., Del Tredici, K., Schultz, C., Thal, D.R., Braak, E. & Braak, H. (2001) The autonomic higher order processing nuclei of the lower brain stem are among the early targets of the Alzheimer’s disease-related cytoskeletal pathology. Acta neuropathologica, 101, 555–564.

Samaey, C., Schreurs, A., Stroobants, S. & Balschun, D. (2019) Early Cognitive and Behavioral Deficits in Mouse Models for Tauopathy and Alzheimer’s Disease. Front Aging Neurosci, 11, 335.

Shimojo, M., Takuwa, H., Takado, Y., Tokunaga, M., Tsukamoto, S., Minatohara, K., Ono, M., Seki, C., Maeda, J., Urushihata, T., Minamihisamatsu, T., Aoki, I., Kawamura, K., Zhang, M.R., Suhara, T., Sahara, N. & Higuchi, M. (2020) Selective Disruption of Inhibitory Synapses Leading to Neuronal Hyperexcitability at an Early Stage of Tau Pathogenesis in a Mouse Model. The Journal of neuroscience : the official journal of the Society for Neuroscience, 40, 3491–3501.

Shrivastava, A.N., Redeker, V., Pieri, L., Bousset, L., Renner, M., Madiona, K., Mailhes-Hamon, C., Coens, A., Buée, L., Hantraye, P., Triller, A. & Melki, R. (2019) Clustering of Tau fibrils impairs the synaptic composition of α3-Na(+)/K(+)-ATPase and AMPA receptors. The EMBO journal, 38.

Smith, W.W., Gorospe, M. & Kusiak, J.W. (2006) Signaling mechanisms underlying Abeta toxicity: potential therapeutic targets for Alzheimer’s disease. CNS & neurological disorders drug targets, 5, 355–361.

Supnet, C. & Bezprozvanny, I. (2010) The dysregulation of intracellular calcium in Alzheimer disease. Cell calcium, 47, 183–189.

Sydow, A., Van der Jeugd, A., Zheng, F., Ahmed, T., Balschun, D., Petrova, O., Drexler, D., Zhou, L., Rune, G., Mandelkow, E., D’Hooge, R., Alzheimer, C. & Mandelkow, E.M. (2011) Tau-induced defects in synaptic plasticity, learning, and memory are reversible in transgenic mice after switching off the toxic Tau mutant. The Journal of neuroscience : the official journal of the Society for Neuroscience, 31, 2511–2525.

Vosler, P.S., Brennan, C.S. & Chen, J. (2008) Calpain-mediated signaling mechanisms in neuronal injury and neurodegeneration. Molecular neurobiology, 38, 78–100.

Waterhouse, B.D., Lin, C.S., Burne, R.A. & Woodward, D.J. (1983) The distribution of neocortical projection neurons in the locus coeruleus. The Journal of comparative neurology, 217, 418–431.

Watt, G., Przybyla, M., Zak, V., van Eersel, J., Ittner, A., Ittner, L.M. & Karl, T. (2020) Novel Behavioural Characteristics of Male Human P301S Mutant Tau Transgenic Mice – A Model for Tauopathy. Neuroscience, 431, 166–175.

Weinshenker, D. (2018) Long Road to Ruin: Noradrenergic Dysfunction in Neurodegenerative Disease. Trends in neurosciences, 41, 211–223.

Williams, J.T., North, R.A., Shefner, S.A., Nishi, S. & Egan, T.M. (1984) Membrane properties of rat locus coeruleus neurones. Neuroscience, 13, 137–156.

Ye, R., O’Callaghan, C., Rua, C., Hezemans, F.H., Holland, N., Malpetti, M., Jones, P.S., Barker, R.A., Williams-Gray, C.H., Robbins, T.W., Passamonti, L. & Rowe, J. (2022) Locus Coeruleus Integrity from 7 T MRI Relates to Apathy and Cognition in Parkinsonian Disorders. Movement disorders : official journal of the Movement Disorder Society, 37, 1663–1672.

Yoshiyama, Y., Higuchi, M., Zhang, B., Huang, S.M., Iwata, N., Saido, T.C., Maeda, J., Suhara, T., Trojanowski, J.Q. & Lee, V.M. (2007) Synapse loss and microglial activation precede tangles in a P301S tauopathy mouse model. Neuron, 53, 337–351.

Zhong, P., Cao, Q. & Yan, Z. (2022) Selective impairment of circuits between prefrontal cortex glutamatergic neurons and basal forebrain cholinergic neurons in a tauopathy mouse model. Cerebral Cortex, 32, 5569–5579.

Zhu, Y., Zhan, G., Fenik, P., Brandes, M., Bell, P., Francois, N., Shulman, K. & Veasey, S. (2018) Chronic Sleep Disruption Advances the Temporal Progression of Tauopathy in P301S Mutant Mice. The Journal of Neuroscience, 38, 10255–10270.

Zweig, R.M., Ross, C.A., Hedreen, J.C., Steele, C., Cardillo, J.E., Whitehouse, P.J., Folstein, M.F. & Price, D.L. (1988) The neuropathology of aminergic nuclei in Alzheimer’s disease. Annals of neurology, 24, 233–242.

